# GIFT – an R package to access the Global Inventory of Floras and Traits

**DOI:** 10.1101/2023.06.27.546704

**Authors:** Pierre Denelle, Patrick Weigelt, Holger Kreft

## Abstract

1. Advancing knowledge of biodiversity requires open-access global databases and workflows. This appears particularly crucial for plants, as ongoing changes pose major threats to this central group of organisms. Having large-scale information on plant distributions, functional traits and evolutionary history will enable the scientific community to improve its understanding of the patterns and drivers of plant diversity on a global scale.
2. The Global Inventory of Floras and Traits (GIFT) is a global database of regional plant checklists that has proven successful in documenting biogeographical and geographical patterns of plants. Since the release of the first version of GIFT, the database kept on expanding. We introduce GIFT version 3.0, which contains 5,169 checklists referring to 3,400 regions. These checklists include a total of 371,148 land plant species, mostly vascular plants, of which 354,848 are accepted species names, and 109 functional traits. This new version uses new resources for taxonomic name standardization, is matched to a new plant phylogeny, comes with a new trait aggregation workflow, and includes additional environmental variables.
3. We also present the `GIFT R`-package, which contains all necessary functions to retrieve distributional, functional, phylogenetic, and environmental data from the GIFT database. The package comes with a dedicated website, https://biogeomacro.github.io/GIFT/, which includes three rich vignettes to guide users in retrieving data from GIFT.
4. The recent development of GIFT and its associated `R`-package provide ecologists with access to one of the largest plant databases. This will foster research into regional to global patterns of plant diversity and their underlying mechanisms. Proper versioning of the database and the ability to retrieve and cite data from any previous and current instance of the GIFT database will ensure the reproducibility of studies that utilize it.

## 1 Introduction

To address the persistent challenges posed by global change, it is crucial to have a comprehensive understanding and documentation of species distributions and other dimensions of biodiversity. While global databases with species ranges and trait information are available for some taxa, such as birds, mammals, amphibians and reptiles (BirdLife International, 2022; Myhrvold et al. 2015), equivalent data for plants still lag behind, although recent efforts have improved their spatial and taxonomic coverage. Together with the Botanical Information and Ecology Network (Enquist et al., 2016), the World Checklist of Vascular Plants (Brown et al., 2023; Govaerts et al., 2021), the Global Naturalized Flora (GloNAF; van Kleunen et al., 2019) and the Global Biodiversity Information Facility (GBIF, www.gbif.org), the Global Inventory of Floras and Traits (GIFT; Weigelt, König, and Kreft 2020) has contributed considerably to improving our knowledge of plant distributions. However, until now, this database of regional plant checklists and species-level traits has not been publicly available. Here, we present the latest update of the GIFT database, with the release of version 3.0, and the `GIFT R`-package which grants open access for users to retrieve data.

### History of GIFT

Knowledge of plant diversity and distributions has grown in parallel with the emergence of global initiatives such as GIFT. GIFT is a global database of regional plant checklists, species level functional traits and environmental region characteristics (GIFT; Weigelt, König, and Kreft 2020). Before launching GIFT, plant diversity research using regional floras was primarily based on species numbers rather than species composition leading to expert-drawn diversity maps (Barthlott et al., 1996; Mutke & Barthlott, 2005), estimated plant richness per ecoregion (Kier et al., 2005) and modelled global species richness (Kreft & Jetz, 2007), the latter being recently updated (Cai, Kreft, Taylor, Denelle, et al., 2022). These efforts highlighted an under-representation of islands which led to global assessments of vascular plants (Kreft et al., 2008; Weigelt & Kreft, 2013) and pteridophyte (Kreft et al., 2010) richness on islands. Islands are particularly well-covered in GIFT, making island biogeography one of the main research areas of papers using GIFT. With checklists available for 1,057 islands larger than 1 km^2^ representing 80.7% of the total island area in the world (Weigelt et al., 2013), GIFT is the most comprehensive plant database for studying patterns of island biogeography. This high coverage has been used to characterize island environmental features (Weigelt et al., 2013; Weigelt & Kreft, 2013) as well as the past and present drivers of plant diversity on islands (Cabral et al., 2014; Schrader et al., 2020; Weigelt et al., 2016).

The main advantage of GIFT compared to the earlier data compilations of regional floras is that it includes not only species numbers but the species composition per region, the floristic status of the occurrences and functional traits at the species level. Recent studies using the GIFT database could therefore provide global assessments for specific plant groups such as epiphytes (Taylor et al., 2022), phylogenetic diversity (Cai, Kreft, Taylor, Schrader, et al., 2022; Weigelt et al., 2015) and endemism (Cai, Kreft, Taylor, Schrader, et al., 2022), and β-diversity (König et al., 2017). Unlike other databases where species composition is available at the plot level (e.g. sPlot, Bruelheide et al., 2019) or comes from an aggregation of georeferenced occurrences (e.g. GBIF, www.gbif.org), regional-level species composition data in GIFT has high geographic coverage and is spatially less biased while being coarser in resolution/grain (König et al., 2019). Functional trait data in GIFT mainly comes from species descriptions in floras, checklists and online databases and is mainly available at the species level, while trait data in other databases like TRY (Kattge et al., 2011) are usually individual measurements making GIFT data a complementary addition. Functional traits in GIFT have been used to map the distribution of growth and life forms on a global scale (Taylor et al., 2023) and to quantify taxonomic and functional disharmonies between island and mainland floras (König et al., 2021; Razanajatovo et al., 2019; Taylor et al., 2019, 2021; Zizka et al., 2022). Finally, GIFT has also been used to advance our understanding of the distribution of alien plant species compared to native species (Bach et al., 2022; Omer et al., 2022; van Kleunen et al., 2015; Yang et al., 2021) as well as species interactions (Delavaux et al., 2019; Luo et al., 2022).

### From GIFT 1.0 to GIFT 3.0

Since the release of version 1.0 in 2018 (https://gift.uni-goettingen.de/about), the number of plant checklists and geographical regions in GIFT has steadily increased (Figure 1). With 5,169 checklists, GIFT 3.0 contains 50% more checklists than version 1.0. These checklists refer to 3,400 georeferenced regions. 2,899 regions have at least one checklist for all native vascular plants (Figure 1a). Different combinations of taxonomic group, floristic status and their completeness can be queried, leading to different numbers (Figure S1). Altogether, these regions cover the entire terrestrial surface of the Earth, as shown on the GIFT website: https://gift.uni-goettingen.de/map (Figure 2a). Across all checklists, GIFT 3.0 includes a total of 371,148 standardized plant species (`work_ID` in the database). These derive from state-of-the-art standardization of 1,161,174 unstandardized original names (after correction of genus names and exclusion of hybrid names; `name_ID` in the database, Figure S2) that differ in spelling, in the availability of author names or infraspecific information. The taxonomic standardization used in GIFT 3.0 is based on the latest taxonomic resources available, and will be updated with future improvements of the reference databases in new versions. 99.1% of all original names could be matched and standardized to an existing species name in the World Checklist of Vascular Plants (Govaerts et al., 2021) or using the TNRS `R`-package (Maitner et al., 2023) and the additional resources considered in there. Synonymy was resolved for 98.2% of all names. In comparison, this percentage was at 90.5% with the previous taxonomic workflow (GIFT 1.0, Weigelt et al., 2020). Only 4.4% of all standardized working names were not considered accepted in the taxonomic resources used in our workflow. 98% of all the standardized working names come with distribution information, meaning that 7,521 plants have only trait data.

**Figure 1.**
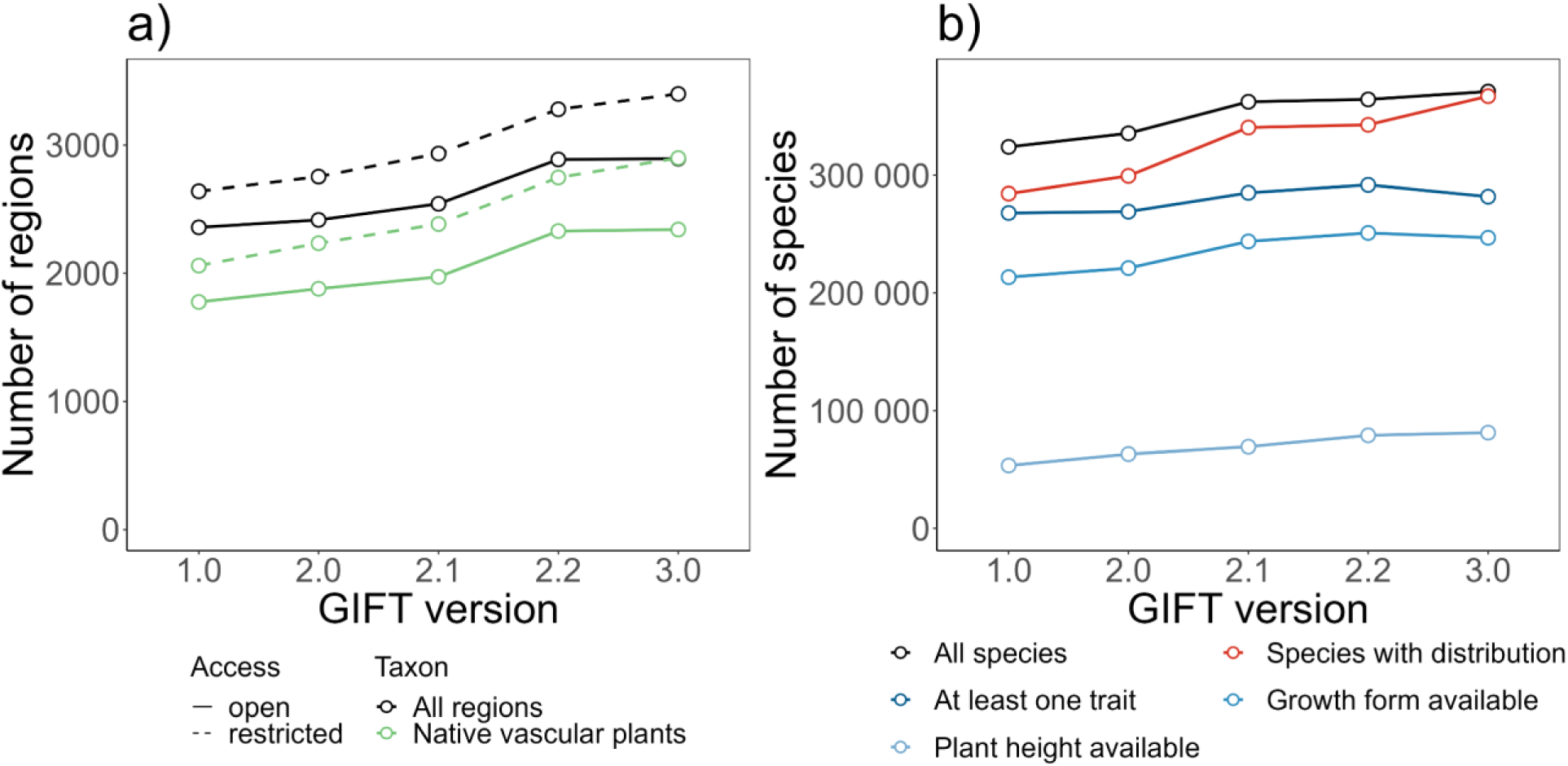
Amount of information in different versions of GIFT. a) Number of regions available for different filters applied. Black lines represent the total number of regions per version, while green lines represent the number of regions with species lists of native vascular plants. Continuous lines represent the information directly available with the `GIFT R`-package and default API settings, dashed lines represent numbers including restricted data which require contacting data owners before using them. b) Number of species with certain trait or distribution information available in the different versions of GIFT. Black color corresponds to the total number of species, red color corresponds to species with distribution information, the three blue colors represent species with at least one available trait, growth form or plant height values. The number of species with growth form available decreases slightly between version 2.2 and version 3.0 due to the change of the taxonomic backbone used to harmonize the species names.

**Figure 2.**
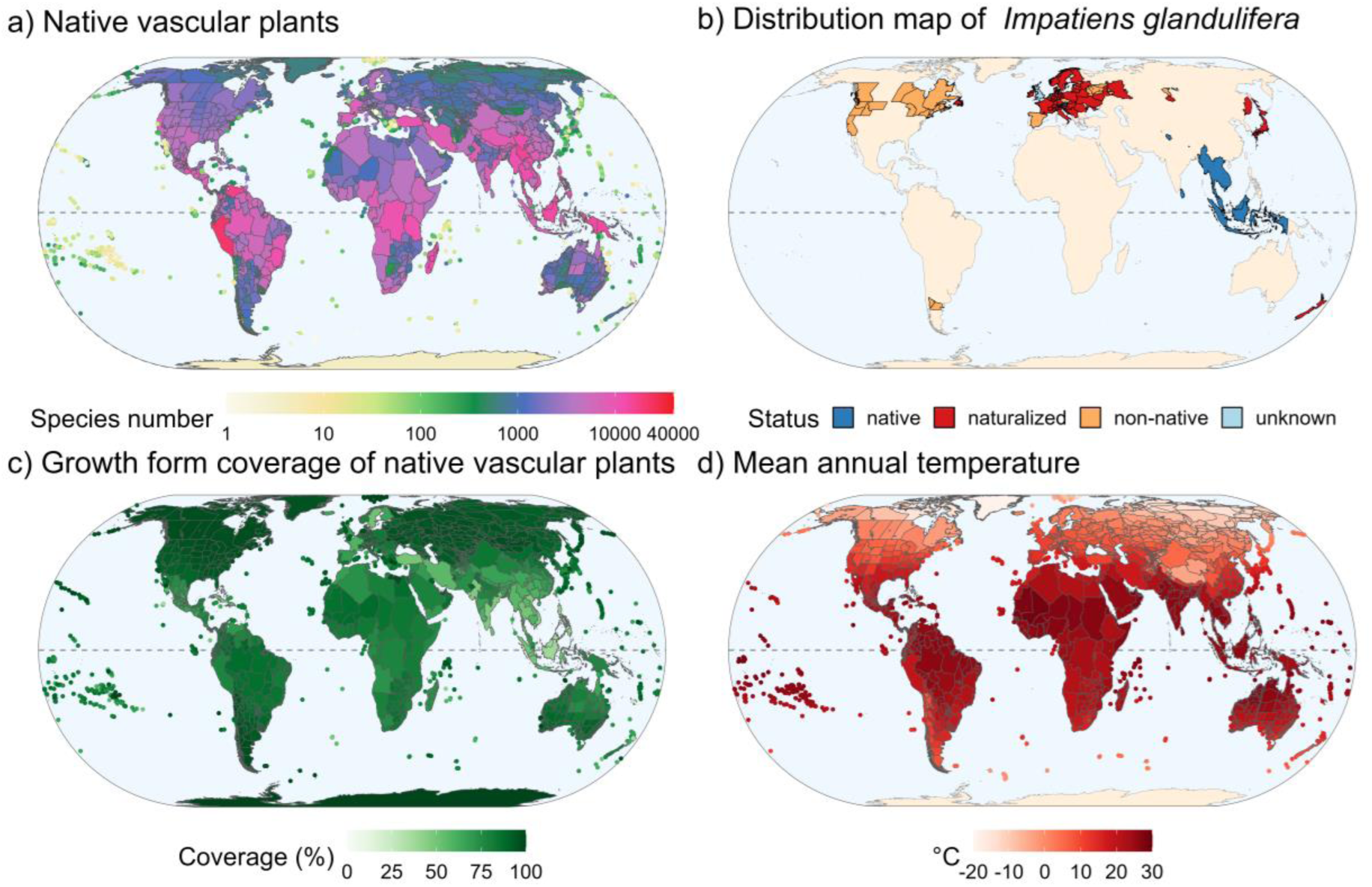
a) Richness of native vascular plants, with a log-transformed color scale, based on `GIFT_richness()`. b) Distribution of *Impatiens glandulifera* with its floristic status in each region as indicated in the primary references used in GIFT, based on `GIFT_species_distribution()`. c) Coverage for the growth form of vascular plants based on `GIFT_coverage()`. d) Mean annual temperature (°C) per region based on `GIFT_env()`. The projection used for all panels is Eckert IV (EPSG: 54012). `R`-code to generate such figures based on functions from the GIFT `R`-package is accessible in the main vignette of the `R`-package https://biogeomacro.github.io/GIFT/articles/GIFT.html. The `GIFT_shapes()` function was used to retrieve the spatial polygons associated with each region.

GIFT 3.0 comes with more trait records than previous versions (Figure 1b, Figure S3), resulting in very high coverage of whole plant traits, such as growth form (246,901 species with growth form) and plant height (81,248 species) which are frequently mentioned in floras and checklists (Figure 1b). 281,836 species have at least one trait available. GIFT 3.0 also uses an updated workflow to aggregate trait information that acknowledges potential disagreement between different references. When querying a categorical trait for a given species, we take the most frequent category as the trait value, indicate alternative trait values and calculate the proportion of references that agree on a given trait value. For continuous traits, the aggregated trait value can be the minimum, mean (e.g. for average plant height) or maximum (e.g. for maximum plant height) of all trait values from the different references included in GIFT. To acknowledge the variation across references, the coefficient of variation is returned.

In GIFT 3.0 all standardized species names are linked to a phylogeny built using the U.PhyloMaker `R`-package (Jin & Qian, 2022) which is stored in the database and accessible via the `GIFT R`-package. For seed plants, it is based on the GBOTB megatree from Smith & Brown (2018) and for pteridophytes, on Zanne et al. (2014), both standardized according to WCVP. Missing species and genera were bound to their respective genera and families using ‘Scenario 3’ from the U.PhyloMaker `R`-package.

To facilitate studies of diversity-environment relationships, GIFT regions are linked to 213 environmental variables. In GIFT 3.0, we added some recently published datasets including paleoclimatic variables (J. L. Brown et al., 2018; Karger et al., 2021), new bioclimatic variables (Brun et al., 2022), global soil temperature variables (Lembrechts et al., 2022), indices of climate stability (Owens & Guralnick, 2019), or aridity (Zomer et al., 2022). We also added the prediction of plant richness per region extracted from the recent work of Cai, Kreft, Taylor, Denelle, et al. (2022). Extraction of raster values for GIFT regions now uses the new `exactextractr R`-package (Baston et al., 2022).

### Package functionality

The `GIFT R`-package contains 27 functions (Table 1) that allow users to retrieve plant checklists matching specific criteria, species distribution information, functional traits, environmental variables, spatial information, and a phylogeny. Most of these functions allow retrieving tables at the global scale or only for a subset of regions or plant species. The `GIFT R`-package comes with a website (https://biogeomacro.github.io/GIFT/), with vignettes, tutorials, and detailed documentation for each function. The `R`-package calls an Application Programming Interface (API), and the API then accesses the database to retrieve the desired data. If necessary, the API can also be called programmatically or through the browser independently of the `R`-package (https://biogeomacro.github.io/GIFT/articles/GIFT_API.html).

**Table 1.** Overview of the 27 functions available in the GIFT R-package.

| Category | Functions | Description |
| --- | --- | --- |
| Regional checklists and species distributions | `GIFT_checklists()` | Checklists for regions matching certain criteria |
|  | `GIFT_checklists_conditional()` | Metadata of checklists for regions matching certain criteria |
|  | `GIFT_checklists_raw()` | Checklists by reference (`ref_ID`) |
|  | `GIFT_lists()` | Metadata of checklists and regions per reference |
|  | `GIFT_species_distribution()` | Distributions of individual plant species |
| Species names, taxonomy and phylogeny | `GIFT_phylogeny()` | Phylogeny linked to standardized species names |
|  | `GIFT_species_lookup()` | Taxonomic information per species name |
|  | `GIFT_species()` | All standardized species available |
|  | `GIFT_taxonomy()` | Taxonomy lookup |
|  | `GIFT_taxgroup()` | Assigns taxonomic groups to species |
| Traits | `GIFT_traits()` | Aggregated trait values per species |
|  | `GIFT_traits_meta()` | Metadata for traits |
|  | `GIFT_traits_raw()` | Non-aggregated reference-level trait values |
|  | `GIFT_traits_tax()` | Trait values at taxonomic group level |
| Spatial and environmental data | `GIFT_env()` | Environmental data per region |
|  | `GIFT_env_meta_misc()` | Metadata for miscellaneous environmental variables |
|  | `GIFT_env_meta_raster()` | Metadata for environmental raster layers |
|  | `GIFT_spatial()` | Regions matching spatial input |
|  | `GIFT_regions()` | Miscellaneous information per region |
|  | `GIFT_shapes()` | Shapefile of selected or all regions |
|  | `GIFT_overlap()` | Spatial overlap between GIFT regions and external polygon resources |
|  | `GIFT_no_overlap()` | Pairwise overlap between GIFT regions |
| **Utils** | `GIFT_coverage()` | Taxonomic and trait coverage per region |
|  | `GIFT_references()` | Metadata per reference |
|  | `GIFT_richness()` | Species richness per region and taxonomic group |
|  | `GIFT_versions()` | Versions of GIFT available |
|  | `western_mediterranean` | Shapefile of the western Mediterranean basin (example data) |

Most of the functions are interconnected and interdependent (Figure S4). Some arguments are shared with other functions, such as the identification numbers for references (`ref_ID`), checklists (`list_ID`), unstandardized species names (`name_ID`), standardized species names (`work_ID`), traits (`trait_ID`) and regions (`entity_ID`). We also provide the possibility to extract and work with any existing version of the database. Each of the functions presented below has an argument called `GIFT_version` that allows retrieving data from a specific instance of the database. Because all versions are available through the `R`-package, analyses can be reproduced even after the database has been updated.

A small number of data providers have asked to restrict access to their data at this time. This implies that among all the data that can be retrieved with the `R`-package, a small proportion of references (6.7%, Figure 1a) are restricted. These data have mostly been contributed by data owners who plan to publish their data independently before making them publicly available via GIFT or come from large online databases that are better accessed directly (e.g. WCVP Brown et al., 2023). A password-protected API, that can be made available upon request, is needed to retrieve these restricted data. On top of the password-restricted API, approval from the data providers is needed. The `GIFT_coverage()` function gives an overview of the potential information available in these references and can be run before considering the use of restricted data. Regardless, we demand citing all data sources used in the references section of studies using GIFT data.

## 2 Regional plant checklists

The main function of the package, `GIFT_checklists(),` allows to retrieve plant checklists for regions that meet certain criteria. A complete tutorial on how to use this function is available in the main vignette of the package, https://biogeomacro.github.io/GIFT/articles/GIFT.html, but in the following, the main options are presented.

First, the function allows to define a taxonomic and a floristic group of interest. The taxonomic group can take several values, e.g. all vascular plants (`taxon_name = “Tracheophyta”`) or other taxonomic groups within land plants (e.g. a plant family, `taxon_name = “Orchidaceae”`). All available options can be viewed by running the `GIFT_taxonomy()` function. The `floristic_group` indicates whether the species retrieved should be native, naturalized, endemic, or if all species should be retrieved. Both arguments have a companion argument, `complete_taxon` and `complete_floristic` respectively, which can be set to `TRUE` or `FALSE`. These arguments define whether only regions that are covered by checklists of the entire taxonomic group or floristic status should be retrieved or if regions covered for only a subset can be returned. For example, with `taxon_name = “Tracheophyta”`, a region covered only by an orchid checklist will only be retrieved if `complete_taxon = FALSE`. Equivalently, with `floristic_group = “native”`, a region covered only by lists of endemic species or only trees will only be retrieved if `complete_floristic = FALSE.` More detailed explanations can be found in the main vignette of the package.

Second, `GIFT_checklists()` works with or without spatial restriction. By default, there is no restriction, meaning that regions covered by checklists meeting the other criteria will be retrieved all over the world. It is also possible to provide a shapefile or a set of coordinates to spatially constrain the query. As an example, we provide a ready-to-use shapefile of the western Mediterranean basin, which can be retrieved using the `western_mediterranean` data function. Regions that fall inside, intersect or encapsulate the spatial input can then be retrieved. Another important consideration is whether users want to retrieve several checklists for nested and overlapping regions or if only non-overlapping regions should be considered. This can be easily set with the `remove_overlap` argument. Overlapping regions can be removed in favor of the smaller or larger regions (above or below a user-defined area threshold) in case of overlap and the removal can be done only within a given reference or overall. If interested in retrieving the species richness of a particular taxonomic and floristic group per region, one can use the alternative function `GIFT_richness()`.

`GIFT_checklists()` returns a list with two objects: the metadata of the regions matching the input criteria, together with the identification numbers of the corresponding references and checklists, and a table with the species composition. By default, the standardized species names and their floristic status are returned in this table, but non-standardized species names can also be returned on request. The original names can help to understand the floristic status assigned to the species, as these statuses refer to the names before taxonomic standardization.

## 3 Distribution of individual plant species

Distribution information from GIFT can also be accessed at the species level. First, to look up if a plant species is included in GIFT, users can call the `GIFT_species_lookup()`function. This function can search for both original or standardized species names and returns all references in which the focal species name occurs as well as the original species name as found in these references. The complete list of standardized plant species names available in the GIFT database can be retrieved with the function `GIFT_species()`. Second, the function `GIFT_species_distribution()`returns a list of regions where the focal plant species occurs. The floristic status of the species is provided, allowing for quick identification of regions where the focal plant is native, naturalized, or endemic. The argument `aggregation` allows to aggregate this information to the region level, with a note indicating whether the original references disagree on the status. An example using *Impatiens glandulifera*, an annual plant native to the Himalayas that is now considered invasive or naturalized in many areas, is shown in Figure 2b). The example illustrates that the floristic status must be treated with caution. For example, while *Impatiens glandulifera* is considered native to the Himalayas, some references also classify it as native to Southeast Asia. This type of conflict, which occurs for many species, may be due to conflicting references, the fact that the floristic status refers to the original species names before taxonomic standardization, or errors in the references. Alternatively, GIFT data can be used via the bRacatus `R` package to estimate a georeferenced occurrence record’s probability of being true or false and its biogeographic status based on a probabilistic framework (Arle et al., 2021).

## 4 Taxonomy

The GIFT database includes original species names as taken from the original references as well as standardized species names and statistics about the standardization process. Several functions in the package, such as `GIFT_checklists()` or `GIFT_species_lookup()`, can return the original species names before standardization and information about the standardization process. It is important to provide this information because the floristic status of species refers to the original species names, not the standardized names. For example, a subspecies may be described as endemic in a reference and standardized to its parent species which is actually considered non-endemic native, resulting in a misleading floristic status for that standardized species. Species names derived from infraspecific taxa or synonyms are therefore highlighted. The harmonization statistics can hence be used to filter for only accepted species (Figure S2), or species that were not subspecies before standardization, etc. Functions that allow retrieving plant checklists, species distributions or raw functional traits, namely `GIFT_checklists()`, `GIFT_species_distribution()` and `GIFT_traits_raw()`, therefore allow to include in their outputs both original and harmonized species names, as well as details on the standardization and the service used. The original and harmonized names come with an identification number (`name_ID` and `work_ID`, respectively) and allow tracing back the taxonomic standardization or to run a preferred alternative taxonomic standardization (Grenié et al., 2023). There are 1,161,174 original species names for 371,148 standardized species names in GIFT 3.0 (Figure S2).

The taxonomy of GIFT assigns species to their respective genera, families and orders up to higher group levels such as Angiosperms, Spermatophytes, etc. It is stored in a table that can be accessed with `GIFT_taxonomy()`. This function returns a table which can be translated into the Newick tree format (Archie et al., 1986) in which each taxon, down to the genus level, is placed between a left and right border. The `GIFT_taxgroup()`function can be used to place any of the accepted plant species names within this taxonomy and to retrieve their higher taxonomic groups (e.g. family, order, or higher group).

## 5 Phylogeny

The phylogeny in GIFT is based on existing megatrees (Jin & Qian, 2022; Smith & Brown, 2018; Zanne et al., 2014), includes all vascular plants from the GIFT database and can be accessed with the function `GIFT_phylogeny()` from version 3.0 onwards. Higher taxonomic groups from the taxonomy of GIFT that form monophyletic groups in the phylogeny can be used to retrieve a phylogeny for a taxonomic subset of the entire phylogeny. The phylogeny can be retrieved as a tree object, which can then be used directly with other `R`-packages such as `phytools` (Revell, 2023) or `ape` (Paradis et al., 2023), or as a table mimicking the Newick format (Archie et al., 1986). The phylogeny, in combination with data from the previously described functions, allows the calculation of phylogenetic diversity metrics at the checklist or region level or the estimation of trait coverage along the phylogeny of vascular plants (Figure 3).

**Figure 3.**
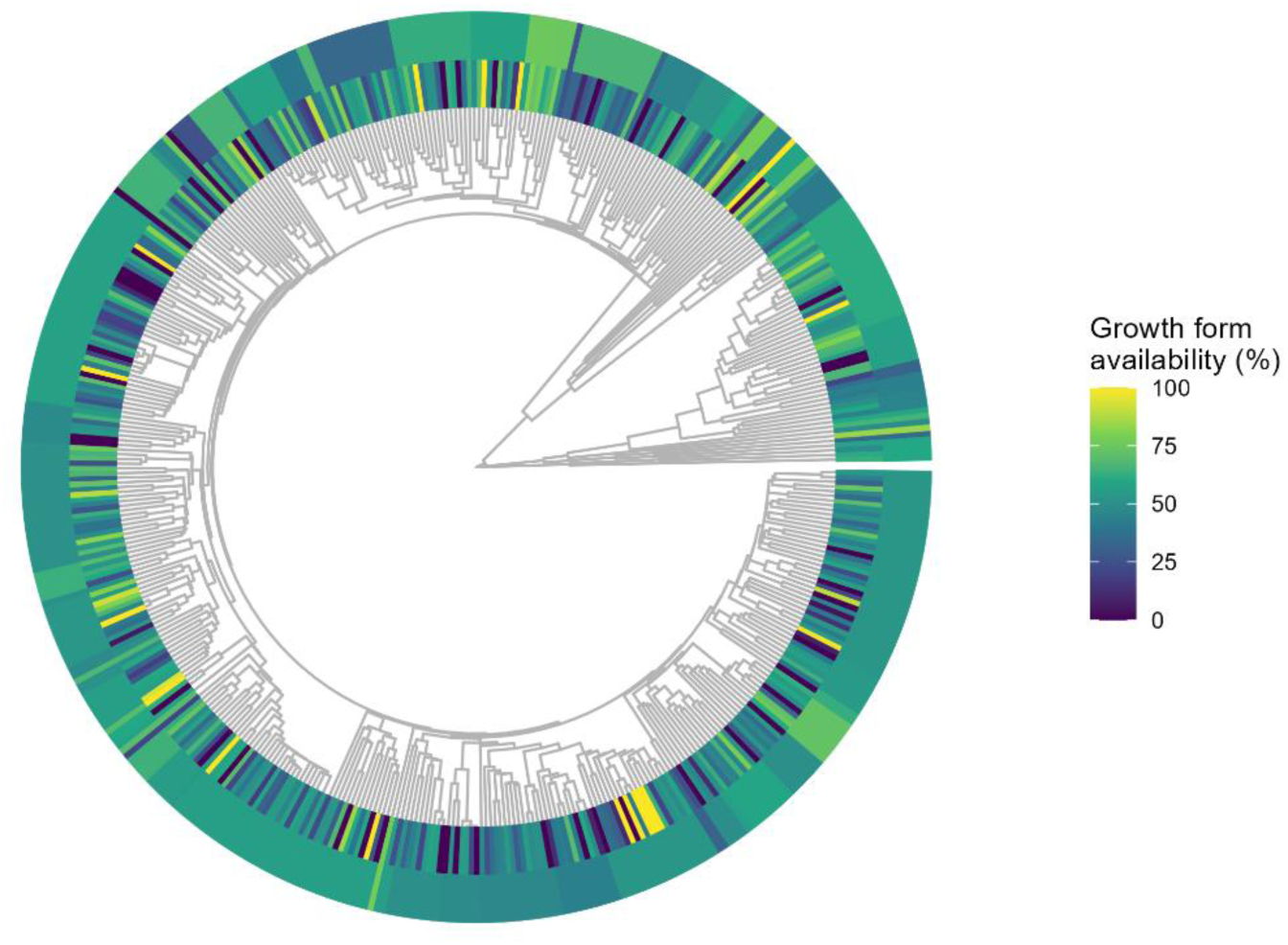
Phylogenetic coverage of the plant trait ‘growth form’. The phylogeny is based on Jin and Qian (2022) and was extracted from GIFT using the `GIFT_phylogeny()` function and pruned to genus level (one tip per genus). The two outer rings illustrate the growth form coverage either at the genus level (inner ring) or at the family level (outer ring). Growth form coverage was calculated as the proportion of species covered per genus and family respectively. The `R-`code to generate this figure is accessible in the main vignette of the `R`-package https://biogeomacro.github.io/GIFT/articles/GIFT.html.

## 6 Functional traits

GIFT 3.0 includes 109 different functional traits and a total of 5,679,014 trait records aggregated at the species level (Figure S3). The different traits are distributed across 6 categories: morphological, ecological, physiological, reproductive, genetic or life history traits (Weigelt et al., 2020). Coverage varies considerably among traits with for example the growth form being available for more than 246,901 plant species while dispersal syndrome is available for only about 9,600 species. Metadata for each of these traits can be viewed by calling the `GIFT_traits_meta()`function. When having a list of traits of interest, users may start by calling this metadata function to retrieve their identification number (`trait_ID`). Once this number is known, users can call the functions `GIFT_traits_raw()` or `GIFT_traits()` to receive one or several traits at a time. The functions return the raw traits or the aggregated trait data at the species level, respectively. In GIFT, there is also trait information collected at the higher taxonomic level. For example, all species belonging to the genus *Abies* are woody. This data can be retrieved using `GIFT_traits_tax()`. Querying the raw traits acknowledges the variation of trait values across references and is therefore of interest to ecologists working at the species level. The primary references for these trait values are also returned. `GIFT_traits()` returns trait values aggregated at the species level. For categorical traits (such as e.g. growth form), the function returns the most frequent trait value, along with an agreement score. This score indicates the percentage of references that leads to this most frequent value out of all references that provide information about this trait. For example, if a plant is described as *tree* in two sources and as *shrub* in a third one, then the value *tree* is returned with an agreement score of 66%. Users can decide on a threshold at which they trust the trait values, or they can include the agreement score and the associated uncertainty in their statistical analysis. For continuous traits, the number of references used to aggregate the trait value is given, together with the coefficient of variation of the continuous value listed in the different references.

Trait records in GIFT can be actual values taken from the references but they can also be logically derived from available information (König et al., 2017). For example, a species can have its growth form described as *tree*, and since all trees are phanerophytes, the life form value of this plant species is set to *phanerophyte*. Whether a trait is derived or not is indicated in the raw trait table returned by `GIFT_traits_raw()`. Trait derivation can introduce biases in the proportional representation of trait categories in assemblages. For example, since all trees are phanerophytes, but not all herbs are therophytes, this derivation leads to an overrepresentation of phanerophytes. Similarly, references of only trees (Beech et al., 2017) or only epiphytes (Zotz et al., 2021) introduce biases. Trait values only based on a reference or a derivation that introduces a bias can be filtered out by setting the `bias_deriv` or `bias_ref` arguments to `FALSE`. More details about the arguments of these two functions can be found in the main vignette of the package (https://biogeomacro.github.io/GIFT/articles/GIFT.html).

Finally, the `GIFT_coverage()`function can be used to map the coverage of a particular trait for a given taxonomic and floristic group. In this case, the argument `what` of the function has to be set to `“trait_coverage”.` Figure 2c) maps the coverage of the maximal vegetative height per plant using this function. This function is complementary to the GIFT website https://gift.uni-goettingen.de/map, where trait coverage can be mapped, and can be used to compare data including or excluding restricted references. In addition, this function can be used to indicate whether a combination of region, taxonomic group and floristic status is covered by at least one checklist for public and restricted data in comparison.

## 7 Environmental variables for a region

A major line of research in macroecology and biogeography is to identify the environmental drivers underpinning species distributions and diversity gradients (McGill, 2019). To facilitate such research, GIFT 3.0. contains 213 variables, distributed across 34 miscellaneous variables and values extracted and aggregated from 179 raster layers. Raster layers mostly contain continuous metrics that provide insights into macroclimatic, topographic, edaphic or paleoclimatic variables or sometimes categorical information like the distribution of soil classes (Hengl et al., 2017). Miscellaneous variables provide information on the geography, geometry, floristic regions, isolation and other characteristics of the GIFT regions based on the region’s spatial shapes and additional feature layers. The two metadata functions `GIFT_env_meta_misc()`and `GIFT_env_meta_raster()` provide an overview of all environmental variables and the underlying data layers. Each of these functions returns the list of available raster layers or miscellaneous variables along with their names and references. Once the names of the variables of interest have been identified, users can call the `GIFT_env()` function for a list of GIFT regions, using their identification number (`entity_ID)`.

When calling a miscellaneous variable, such as the area or the biome that encapsulates the region of interest, a single value is returned. For raster layers, however, users need to provide a set of summary statistics. Indeed, as the raster layers have a finer resolution than most of the GIFT regions, the environmental information needs to be aggregated. For example, one can ask for the average temperature of the raster cells falling into a GIFT region (`GIFT_env(rasterlayer = “wc2.0_bio_30s_01”, sumstat = “mean”)`, Figure 2d), or for the standard deviation of the precipitation values (`GIFT_env(rasterlayer = “wc2.0_bio_30s_12”, sumstat = “sd”)`). The desired summary statistics must be passed to the `GIFT_env()` function using the `sumstat` argument.

## 8 Outlook

This paper introduces `GIFT`, an `R`-package that provides easy-to-use functions for accessing the GIFT database. This tool will support a range of new studies on plant biogeography while allowing for their reproducibility. In parallel, we continue integrating new plant checklists into the GIFT database to increase the coverage of underrepresented regions or environments of the world, to bridge the gap between local and regional studies of plant diversity (Puglielli & Pärtel, 2023), and to increase the coverage of functional traits. Additions will be made to the beta version of GIFT, which will be turned into a new stable version once enough additions have been made and checked for consistency. Updates to the package are submitted to the CRAN repository and archived on the Zenodo repository, with each new release having its own DOI (version 1.1.0 of the package is archived at https://zenodo.org/record/8087130).

## Authorship guidelines

Publications using the GIFT database should cite all primary sources as well as the version of GIFT and the original publication of the GIFT database (Weigelt et al., 2020). All primary references in GIFT are available with the `GIFT_references()` function. If GIFT data are retrieved through the `R`-package, this paper should also be cited. We also encourage authors to provide the `R`-scripts used to retrieve the data as supplementary files to their study to promote traceability and reproducibility. A few data contributors of GIFT asked to restrict access to their data for now. These data can be accessed through the `R-`package specifying a password-protected API (provided by the authors of this article upon request) and their use requires contacting the authors of the primary restricted data and asking for permission to use them. We encourage researchers using GIFT to contact us if any help is needed regarding the use of GIFT data.

## Acknowledgments

We thank Martin Turjak, Matthias Grenié and other beta-testers of the `R`-package for providing useful insights. HK acknowledges funding from the DFG as part of the research unit FOR 2716 DynaCom and research training group RTG 1644 Scaling Problems in Statistics.

## Author’s contributions

PD and PW developed the `GIFT R`-package and the associated GitHub website. PW led the development of the database infrastructure and API. All authors worked on the advancement of the GIFT database. PD led the writing of the manuscript to which all authors contributed critically.

## Data accessibility

The `GIFT R`-package is available for download from CRAN at https://CRAN.R-project.org/package=GIFT. The development version of the package is available at https://github.com/BioGeoMacro/GIFT. An associated website with tutorials is available at https://biogeomacro.github.io/GIFT/ and an interactive overview of the data available can be found at https://gift.uni-goettingen.de.

## Supporting information

**Figure S1.**
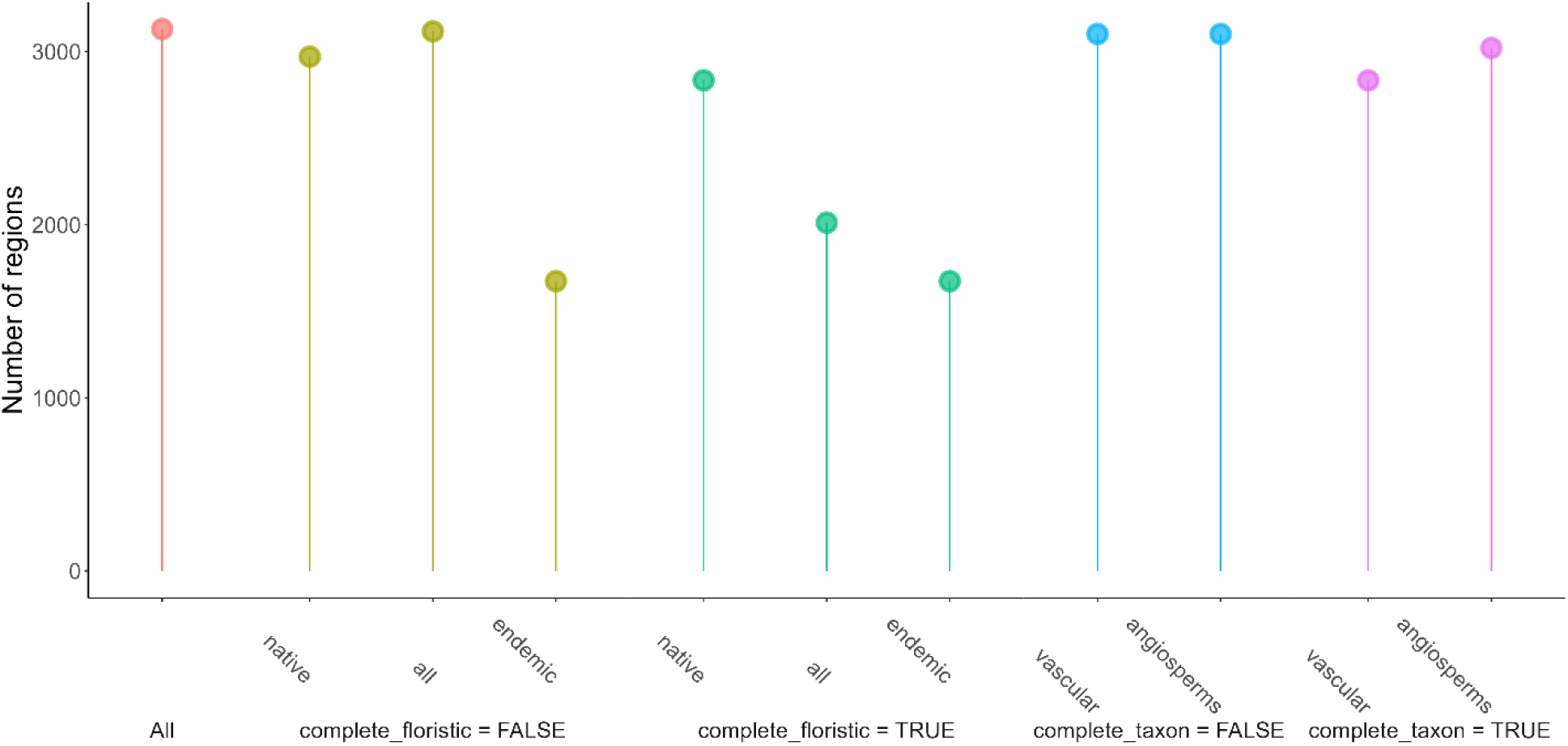
Number of regions retrieved with `GIFT_checklists()` using different criteria for GIFT 3.0. The first red lollipop corresponds to the black dashed line in Figure 1a. For the yellow and green lollipops, the taxonomic group is set to Tracheophyta, i.e. vascular plants and `complete_taxon` is set to `TRUE`. With `complete_floristic = TRUE`, the number of regions with the floristic status set to native is higher than with the floristic status set to all because it is easier to get a complete knowledge of only native species in a region than for all species. For the blue and purple lollipops, the floristic status is set to native and `complete_floristic` is set to `TRUE`. With `complete_taxon = TRUE`, the number of regions with angiosperms is higher than with vascular plants because it is easier to get a complete knowledge of only angiosperms in a region than for all vascular plants.

**Figure S2.**
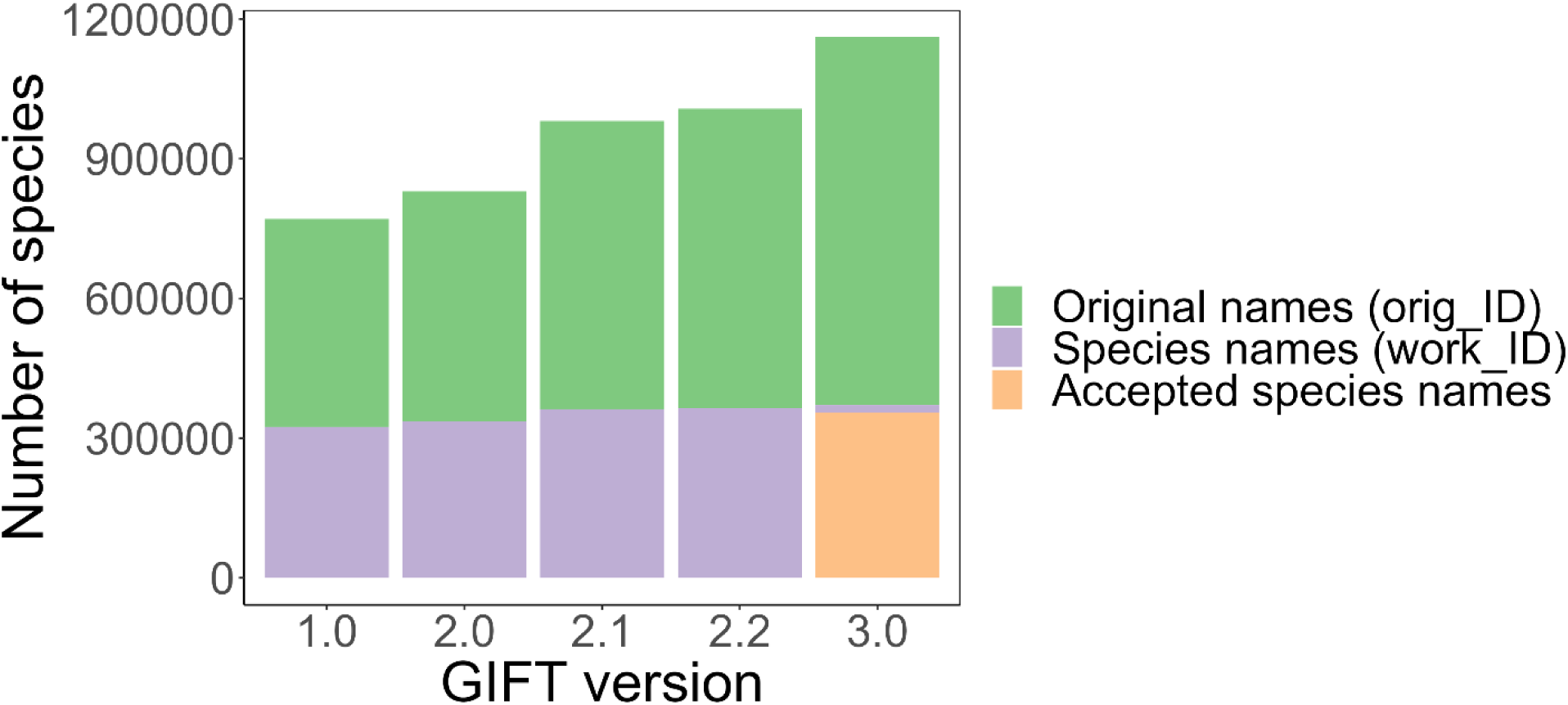
Number of original unstandardized (green) and standardized (purple) species names compared across versions of GIFT. In GIFT 3.0, the new taxonomic workflow using the World Checklist of Vascular Plants (Govaerts et al., 2021) allows to classify species as accepted or not. The majority (95.6%) of standardized names are considered as accepted.

**Figure S3.**
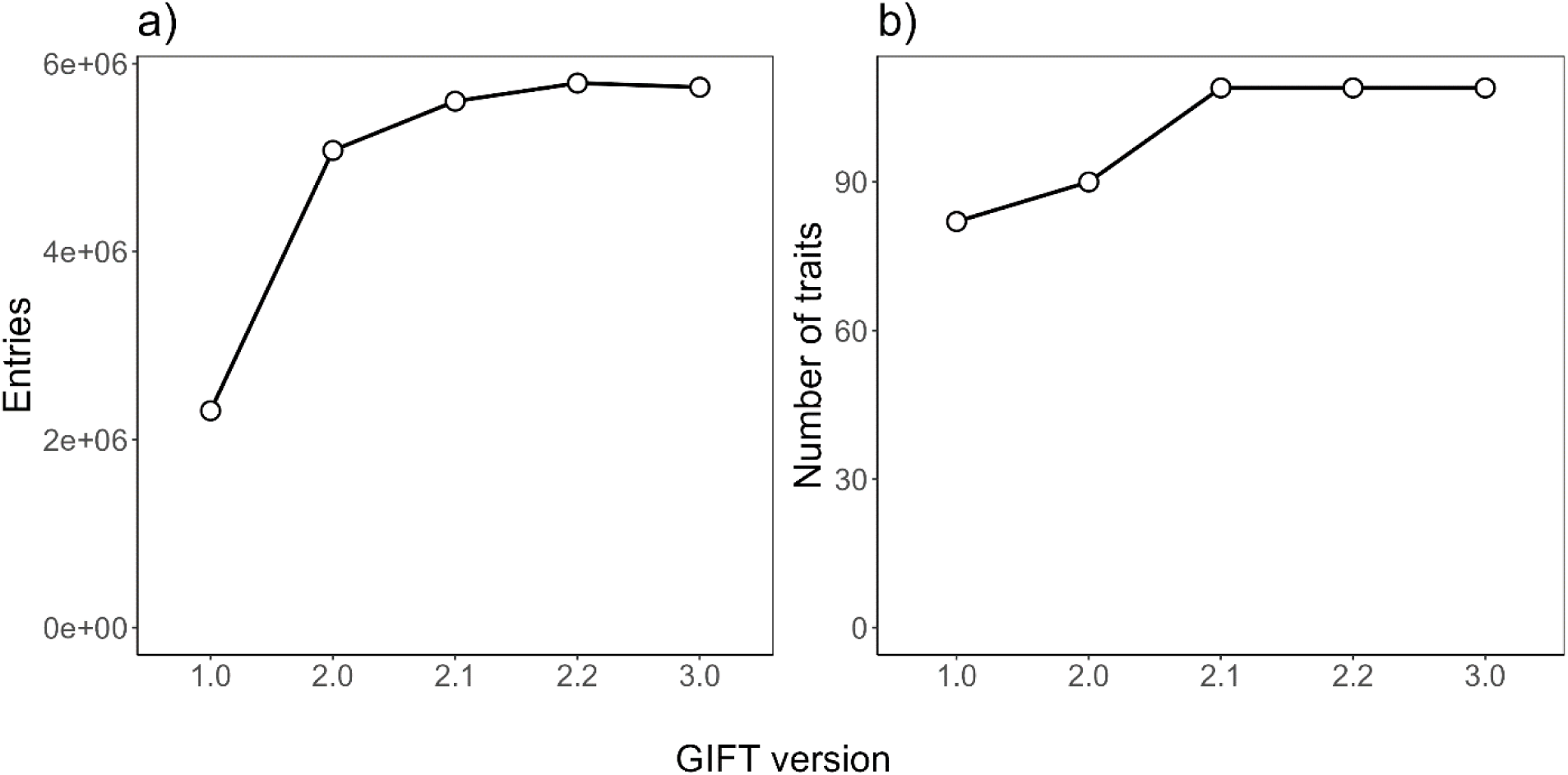
Number of trait records, and number of traits available in the different versions of GIFT. Panel a) indicates how many trait values are available (a record is a value of a particular trait at the species level) and panel b) shows the total number of traits.

**Figure S4.**
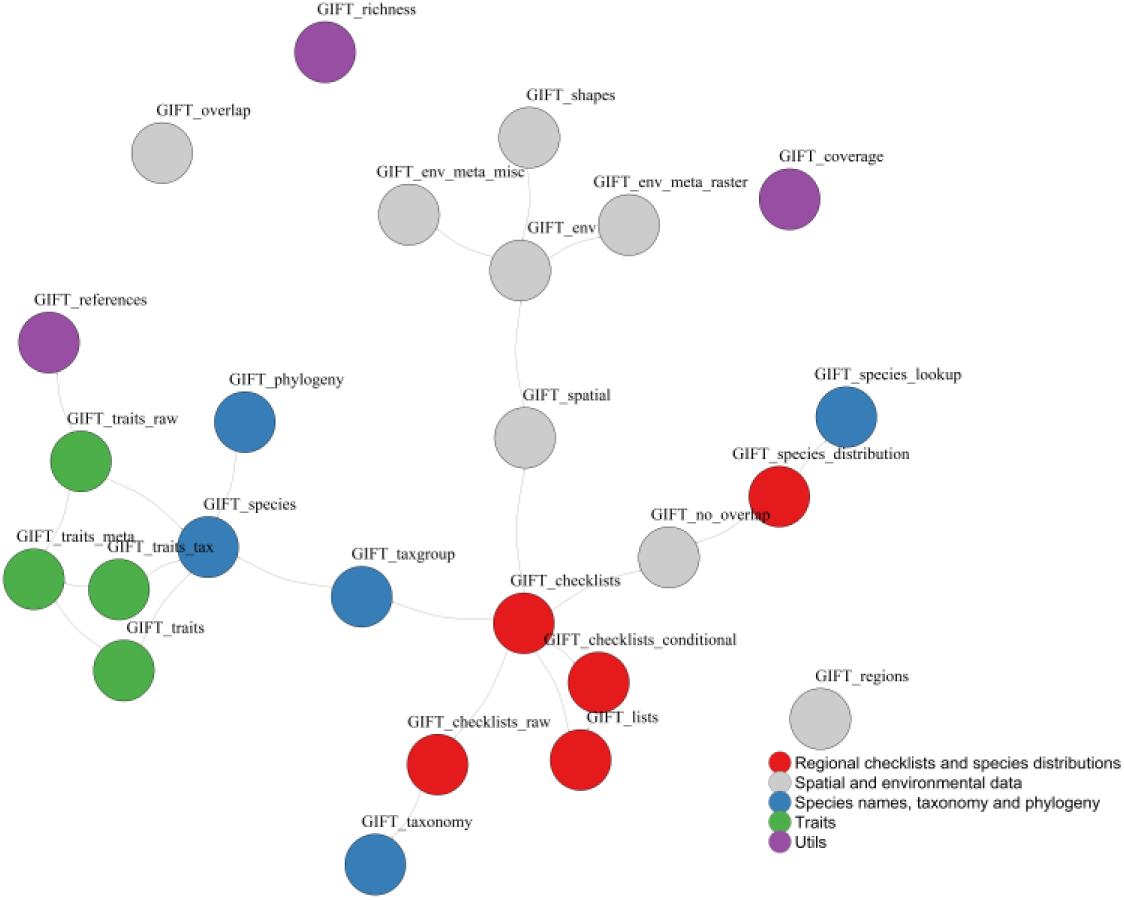
Dependency graph of the 27 functions in the `GIFT R` package. Each circle represents one function. If one function uses another one, then it is connected to it with a grey arrow. Functions are colored according to their category (Table 1). The metadata function `GIFT_versions()` is removed from this graph because it is a metadata function that connected to all other functions.

